# Panomap: Unbiased Nanopore Signal Mapping with Pangenome Variation Graphs

**DOI:** 10.64898/2026.07.10.737796

**Authors:** Po Jui Shih, Zephan Sanghani, Andrea Guarracino, Hasindu Gamaarachchi, Christopher Batten

## Abstract

**Motivation:** Signal-space nanopore mappers enable real-time mapping and filtering decisions directly from raw nanopore signals. However, existing signal-space mappers are built around linear references, and using a single representative reference can introduce reference bias when the sample diverges from that reference. Pangenome reference collections can reduce this bias by representing diversity across related reference sequences, but linear-reference signal mappers must treat each sequence as a separate target, redundantly storing shared sequences. Pangenome variation graphs provide a more compact representation by storing shared sequences once and encoding variants as alternative paths through the graph. Although sequence-to-graph mapping is well established for basecalled reads, existing signal-space methods do not directly use pangenome variation graphs.

**Results:** We present Panomap, the first signal-space mapper that operates on pangenome variation graphs. Panomap maps raw nanopore signals to graph references, allowing signal-space mapping to use pangenome diversity while representing shared sequences once. We evaluate Panomap in three settings. First, when a single reference already maps the sample well, Panomap preserves mapping accuracy as additional reference sequences are added to the reference collection, while state-of-the-art signal-space tools regress. Second, when the exact sample strain is absent from the reference collection, Panomap benefits from adding related assemblies from the same species to the pangenome reference. Third, using a highly polymorphic locus, we show that Panomap can map reads from alleles not represented in the reference collection by using related alleles in the pangenome, with the largest gains for more divergent alleles and for decisions made from short prefixes of the read signal. In addition, Panomap’s graph index scales sublinearly with pangenome collection size. Together, these results show that Panomap brings population-aware reference representation into signal-space mapping.

**Availability and Implementation:** Panomap is open source and available at https://github.com/cornell-brg/panomap.

## Introduction

Nanopore sequencers produce long reads by passing DNA through a protein pore and measuring changes in ionic current as the molecule moves through the pore [1]. Unlike other platforms, nanopore sequencers stream this raw signal to the host computer while sequencing is still ongoing [2]. This enables real-time analysis in two distinct ways. First, analysis can begin before sequencing completes, overlapping computation with data generation and reducing time to result. Second, in targeted sequencing, the analysis can feed decisions back to the sequencer to reject reads that are unlikely to be of interest, a process known as adaptive sampling [3–9].

Several tools now analyze nanopore signals directly, making mapping or filtering decisions from raw signal rather than from basecalled reads. This can reduce the computation needed for early decisions and allow signal-space methods to act as upstream filters for downstream analysis. UN-CALLED [4] maps raw signal to a reference using FM-index-based probabilistic event matching. Sigmap [10] introduced a streaming signal-space mapper using k-d-tree indexing, seed selection, and colinear chaining. RawHash [11] and RawHash2 [2] use hash-based seed-and-chain mapping over quantized signal tokens. Signal-space analysis also extends beyond mapping. Raw nanopore signals contain information that is lost in the basecalled sequence: base modifications such as 5-methylcytosine alter the ionic current, so evidence for these modifications is present in the raw signal rather than in the called bases. Nanopolish [12], Tombo [13], f5c [14], and Uncalled4 [15] align raw signal to a reference to detect these modifications. Rawsamble [16] performs de novo assembly directly from raw nanopore signals. Beyond these software methods, hardware-accelerated systems such as SquiggleFilter [17] and HARU [18] show that direct signal analysis can support selective sequencing on resource-constrained devices.

Across these signal-space tools, a common operation is matching reads to a reference. The choice of reference therefore matters. A single linear reference represents only one sequence and cannot capture the diversity present across a population, from single-nucleotide differences to large structural rearrangements. Pangenomes address this limitation by representing diversity across multiple related reference sequences. Specifically, pangenome variation graphs provide a compact representation of this diversity, storing shared sequences once and encoding variants as alternative paths through the graph. The graph structure can also encode within-genome repeats as cycles [19, 20]. These graphs have been widely adopted in base-space mapping [21–24].

Signal-space tools are built around linear references, so a pangenome can be supplied to them only as a collection of individual reference sequences, not as a single graph. Seed-and-chain mappers such as RawHash2 [2] can index these sequences as a multi-FASTA, treating each as a separate target. As a result, shared sequences are stored redundantly, and a read matching several related reference sequences yields separate mapping results. Sigmoni [25] also operates on a collection of linear references, and its compressed index scales better to large pangenome collections. However, for both RawHash2 and Sigmoni, the input reference remains a set of independent linear sequences, without representing how those sequences relate to one another. A pangenome variation graph encodes both, storing shared sequences once and capturing their relationships in its topology, yet no signal-space tool takes such a graph as input.

We present Panomap, the first signal-space mapper that operates directly on pangenome variation graphs. Panomap extends hash-based seed-and-chain signal mapping to graph references by indexing the paths embedded in the graph, chaining anchors within each path, and merging chains that map to the same shared graph region, all in signal space. This lets Panomap index shared sequences once, then relate mapping results across paths in the graph instead of reporting them only as independent linear-reference hits.

Our results show that Panomap enables signal-space mapping to use pangenome diversity without duplicating sequences shared across separate linear references. We demonstrate this through three evaluations. First, when a single reference already maps the sample well, Panomap maintains mapping accuracy as the pangenome collection grows, while state-of-the-art signal-space tools regress. Second, when the exact sample strain is absent from the reference collection, Panomap improves mapping by adding related assemblies of the same species to the pangenome. Third, at a highly polymorphic locus, Panomap maps reads from alleles not represented in the reference by using related alleles in the pangenome, with the largest gains for more divergent alleles and for decisions made from read signal of limited lengths. In addition, Panomap’s index scales sublinearly with pangenome collection size. Together, these results show that Panomap brings population-aware reference representation into signal-space mapping.

## Methods

Figure 1 shows the Panomap workflow, including the external graph-building step that produces the reference graph in the GFA format (Figure 1a). Panomap is implemented in C++20 and takes this graph and raw nanopore reads in BLOW5 format [26] as input. After processing each read, Panomap emits mappings in PAF or Graph Alignment Format (GAF) [23]. At a high level, Panomap follows two phases: indexing the graph reference and mapping raw nanopore reads.

**Fig. 1.**
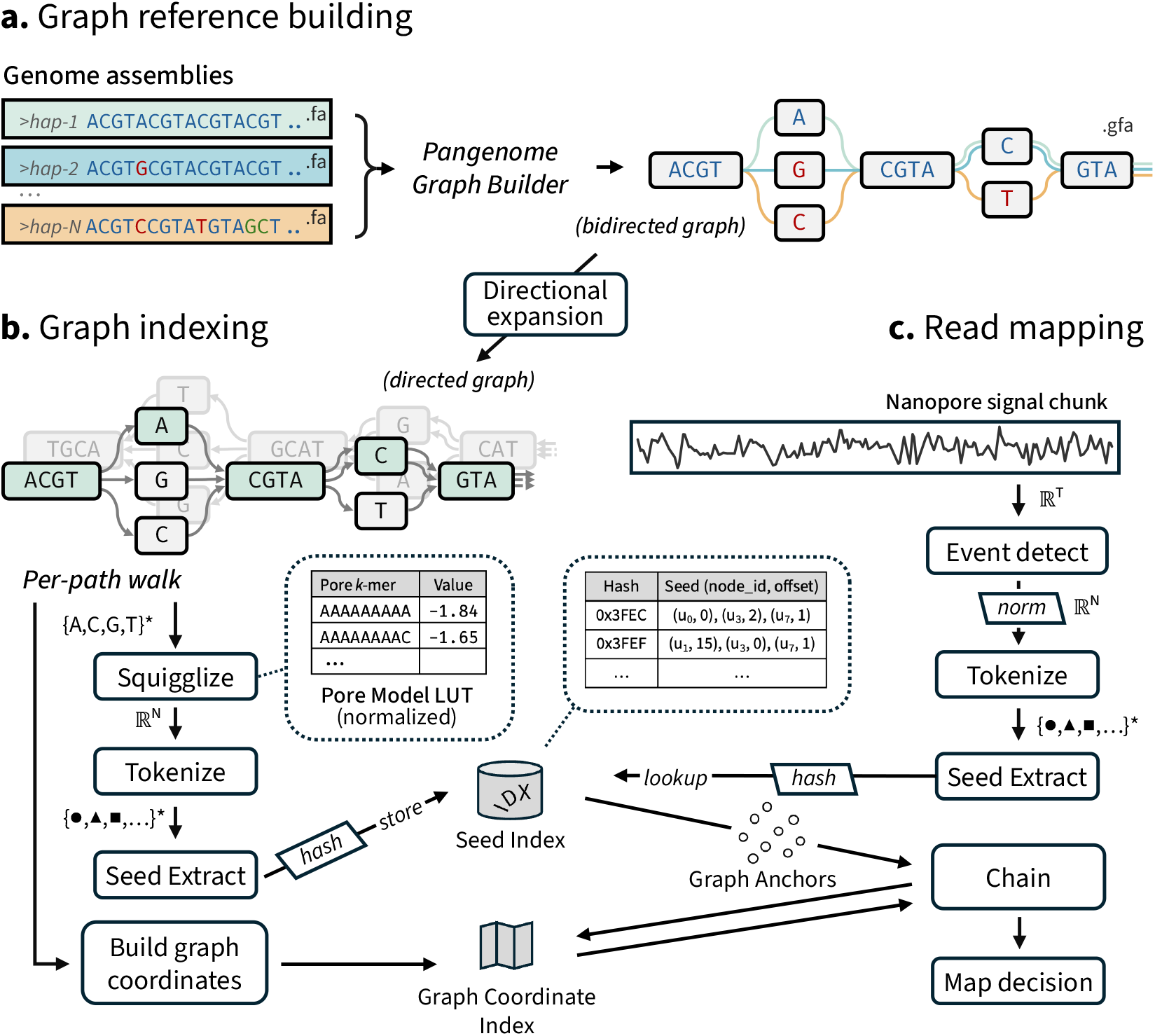
Panomap overview, showing (a) graph reference building, (b) graph indexing, and (c) read mapping.

During indexing, Panomap expands the bidirected input graph into a directed graph and walks the paths embedded in the graph (Figure 1b). From these paths it builds two indexes. The first is a seed index, which maps hashed token seeds to the graph locations where they occur. Panomap builds it by *squigglizing* each path’s base sequence into its expected event stream, *tokenizing* the events into discrete symbols, and hashing the result into seeds. The second is a graph coordinate index that records where each node visit falls so Panomap can measure anchor distances along each path and detect chains that overlap across paths.

During mapping, Panomap turns each read’s raw signal into tokens in the same symbol space used during indexing, so read seeds can match the indexed reference seeds (Figure 1c). It *detects* events by segmenting the signal into regions of similar current, *tokenizes* those events, looks up exact seed matches in the index, *chains* the resulting anchors, and emits a mapping decision.

### Pangenome graph input and directional expansion

Panomap takes a pangenome variation graph in GFA format as the reference input. This graph is typically constructed from a collection of input reference sequences using tools such as pggb [27] or Minigraph-Cactus [28] (Figure 1a). pggb constructs the graph reference-free through all-to-all alignment of input sequences (`wfmash` [29]) followed by graph induction (`seqwish` [30]), whereas Minigraph-Cactus starts from a primary reference genome and iteratively adds structural and small variants from the remaining input sequences. Both approaches produce graphs with nodes (nucleotide sequences), edges (adjacencies between nodes), and paths (ordered walks over nodes from input sequences). In these graphs, sequences shared across input sequences appears once as shared nodes, while sequence differences appear as alternative paths through the graph. Path information is essential for Panomap’s per-path indexing and chaining, so graph representations without paths, such as Minigraph’s output [23] or Bifrost’s colored compacted de Bruijn graph [31], are not supported.

The input GFA is a bidirected sequence graph: each node can be traversed in either the forward or reverse-complement orientation. These two traversals correspond to different nucleotide strings and therefore produce different expected event streams under the pore model. As shown by the *directional expansion* step in Figure 1, Panomap converts the bidirected input graph into a directed graph by creating separate forward and reverse-complement copies of each node, edge, and path. This makes strand orientation explicit before indexing. Rather than indexing one graph and handling reverse-complement matches as a separate case during mapping, Panomap indexes directed graph paths whose nucleotide sequences are already orientation-specific.

### Per-path signal seed indexing

As part of graph indexing (Figure 1b), Panomap builds the seed index by first z-score normalizing the pore-model table used for squigglization. The pore-model table defines the expected nanopore event value for each nucleotide *k*_pore_-mer. Panomap then walks each path through the directed graph. For each path, a window of width *k*_pore_ slides over the path sequence, and each resulting *k*_pore_-mer is looked up in the normalized poremodel table. The sequence of lookup results forms the path’s expected event stream. An event-level homopolymer compression (HPC) step then merges consecutive events whose normalized values differ by less than a small threshold, analogous to base-space HPC in minimap2 [32].

The resulting event stream is tokenized with the 4-bit adaptive quantizer used by RawHash2 [2], which maps each normalized event value to one of 16 discrete tokens. Panomap then extracts seeds by hashing each window of *k*_seed_ consecutive tokens. Each seed is stored in the seed index as a seed record (*u, o*) under the token-seed hash *h*, where *u* is the graph node containing the seed and *o* is the seed’s start offset within node *u*.

Both squigglization and seed extraction require multi-base context. Parsing each node sequence alone misses context at node ends, while enumerating all possible node-edge walks through variant forks can quickly become impractical. Per-path walking avoids this problem by indexing only the paths stored in the graph. The seed index maps each seed hash *h* to a list of graph locations where that seed occurs. Because different paths can visit the same shared node, path walking may emit the same graph location (*u, o*) more than once for the same seed hash. After path walking, Panomap deduplicates seed records so that each graph location (*u, o*) appears at most once for each seed hash *h*. During mapping, Panomap looks up each read seed in the seed index and retrieves a list of unique graph locations for that seed. By removing duplicate records from shared graph sequence, this deduplication lets the seed index scale sublinearly with pangenome size.

### Graph coordinate index

In addition to the seed index, Panomap also builds a graph coordinate index during graph indexing (Figure 1b). This index contains two coordinate structures over the directed graph. The first structure records a path coordinate for each node visit, used for chaining. During the path walk, Panomap tags each node visit with a path coordinate (*p, r*), where *p* is the path identifier and *r* is the cumulative base position along path *p* at the start of the visit. A node may be visited multiple times within the same path or by multiple different paths, so a graph node can be associated with multiple path coordinates. During mapping, these coordinates are used to project graph locations onto the paths that contain them, allowing Panomap to measure anchor distances along each path for colinear chaining.

Panomap stores one path coordinate for each visit, so the same graph node can correspond to multiple path positions. Each path therefore defines an independent linear coordinate system in which anchor distances reflect the true genomic distance along that path. During mapping, these coordinates are used to expand graph locations into positions along each path for colinear chaining.

The second is an approximate global 1D coordinate system over the directed graph, used for chain deduplication. Per-path chaining can produce near-duplicate chains when multiple paths share the same region. To identify these overlaps, Panomap adapts the 1D path-guided stochastic gradient descent (1D PG-SGD) layout algorithm from ODGI [33, 34]. 1D PG-SGD places graph nodes along a line so that distances between nodes roughly agree with their nucleotide distances along the graph paths. Panomap modifies ODGI’s 1D PG-SGD in two ways. First, each node receives start and end coordinates rather than ODGI’s single start position, so intra-node positions can be interpolated and chains can be projected as 1D intervals. Second, the graph is partitioned into connected components, and coordinates restart from zero within each component, since disconnected components share no distance along any path. These per-node coordinates are later used to detect overlapping chains.

### Signal processing and anchor formation during mapping

During read mapping (Figure 1c), Panomap maps each read incrementally, processing one signal chunk (a fixed-size block of raw signal samples) at a time. Within each chunk, the signal passes through a signal-space preprocessing pipeline that converts the raw signal to picoampere (pA) current, detects events using Scrappie’s two-window t-statistic detector [35], normalizes each chunk by z-score, and applies the same event-level HPC used during indexing. The event stream is then tokenized with the same adaptive quantizer used during indexing, and seeds are extracted by hashing windows of *k*_seed_ consecutive tokens. Each seed hash is queried against the seed index, returning matching graph locations (*u, o*). Combined with the seed’s read position *q*, each match forms a *graph anchor* (*q, u, o*).

### Per-path colinear chaining

From the previous step, Panomap obtains a set of graph anchors (*q, u, o*). Using the path coordinates from the graph coordinate index, Panomap expands each graph anchor into one *path anchor* (*q, r*) for each path that visits its node, where *r* is the reference position on that path. It groups these path anchors by path and chains them independently within each path using the colinear chaining formulation of RawHash2, which is based on minimap2 [32]. This procedure rewards anchors that progress consistently along both the read and the path, while penalizing large or uneven gaps, producing high-scoring chains as candidate mapping locations.

Panomap identifies chains in decreasing order of chain score, reconstructing each by traceback from its endpoint anchor. It keeps chains that meet minimum score and anchor-count thresholds and whose scores are at least a configurable fraction of the top chain score. Panomap also deduplicates chains using the global 1D coordinates from the graph coordinate index. Each chain’s start and end graph anchors are projected onto these coordinates, and a chain whose projected interval overlaps a higher-scoring chain in the same connected component above a configurable threshold is discarded as a duplicate from the same region. This leaves one representative chain per genomic region.

### Mapping decision

After each chunk, Panomap evaluates the top candidate chains and decides whether to emit a mapping. It applies one of two rules depending on whether multiple chains compete or only one chain is found. With multiple chains, Panomap computes a weighted decision score adapted from RawHash2 [2]. This score is high when the best chain has high mapping quality and is clearly separated from the other chains in both mapping quality and chain score. Let *Q*_best_ and *C*_best_ be the mapping quality and chain score of the best chain, and let 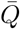 and 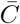 be the corresponding means across competing chains. Defining *r_q_* = min(*Q*_best_ /30, 1), 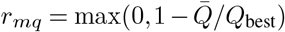, and 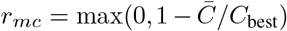, the weighted decision score is *w*_sum_ = *w*_*q*_*r*_*q*_ + *w*_*mq*_*r*_*mq*_ + *w*_*mc*_*r*_*mc*_, where *w*_*q*_, *w*_*mq*_, and *w*_*mc*_ are configurable weights. Panomap emits a mapping if *w*_sum_ exceeds a configurable threshold. The chain deduplication described above provides this rule with a non-redundant chain set; otherwise, near-duplicate chains across paths would make the best chain appear less distinct from its competitors.

When only one chain is extracted, the weighted decision rule does not apply. Panomap accepts the single chain if its anchor count exceeds a minimum threshold and its read and reference event spans are comparable under a configurable span-ratio threshold. If no mapping is emitted for the current chunk, Panomap retains the top candidate chains’ anchors and processes them together with the next chunk. This chunk-by-chunk process continues until Panomap makes a confident decision or reaches *c*, the configurable maximum number of signal chunks it observes before deciding. A read still undecided at *c* is reported as unmapped. Deciding as early as possible reduces per-read processing time. Mappings are emitted in PAF or GAF format with per-read tags.

## Results

The results are organized into four subsections. The first three evaluate Panomap’s mapping accuracy across different settings, and the fourth evaluates the cost of using pangenome references in terms of index size and per-read processing time. All experiments ran on an Intel Xeon Gold 6240 server with 72 physical cores, 144 threads, 376 GB RAM, and RHEL 8.10. Each tool uses 64 threads for mapping, with per-tool commands provided in the supplementary material.

### Preserving accuracy under pangenome scaling when one reference already represents the sample

We first consider a setting where reads are already well represented by one reference. In this case, a pangenome mapper should maintain its accuracy as the reference collection grows.

We evaluate Panomap against RawHash2 and Sigmoni using two binary classification tasks. The first task asks whether each tool can keep SARS-CoV-2 reads while rejecting unrelated *E. coli* reads. The second task uses the same setup for *S. cerevisiae*, asking whether each tool can keep *S. cerevisiae* reads while rejecting unrelated *E. coli* reads. The on-target reads are real nanopore reads: 40,000 SARS-CoV-2 reads from InterARTIC [36] and 40,000 *S. cerevisiae* reads from SRA `SRR8648503`. The unrelated off-target reads for both classification tasks are 40,000 simulated *E. coli* reads generated with Squigulator [37] using the R9.4.1 preset from the *E. coli* K-12 MG1655 assembly (NCBI `NC_000913.3`). For both binary classification tasks, we scale the target reference collection from 1 to 8 reference sequences (full composition in Supplementary material Table S2). We write *N* for the number of target reference sequences in the collection, sampled at *N* = 1, 2, 4, 8, and *c* for the maximum number of signal chunks a tool sees before making a decision. Each chunk is configured to be 4000 signal samples. We report recall, specificity, and *F*_1_, where recall is the fraction of on-target reads kept and specificity is the fraction of off-target reads rejected. Panomap maps to pangenome graphs built with pggb and Minigraph-Cactus (MC). RawHash2 and Sigmoni are given the same target pangenome collection, but represented as multi-FASTA references rather than as a graph. Sigmoni additionally includes a fixed *E. coli* reference as the off-target reference required by its binary classification mode.

From Figures 2a and 2b, Panomap maintains its accuracy as the collection grows. Across all four collection sizes (*N* = 1, 2, 4, 8), its *F*_1_ stays near 0.94 on SARS-CoV-2 and 0.89 on *S. cerevisiae* at *c* = 8, and the pggb and MC graphs behave the same. In contrast, RawHash2 and Sigmoni regress as the reference collection scales. RawHash2 is strong at *N* = 1, though on SARS-CoV-2 its specificity worsens as the maximum number of signal chunks *c* grows, reaching 0.07 at *c* = 8. As more reference sequences are added, RawHash2’s recall falls, most sharply at small *c*: one-chunk recall on *S. cerevisiae* drops from 0.43 at *N* = 1 to 0.04 at *N* = 2, lowering its *F*_1_. RawHash2 treats each reference sequence as a separate target, so reads from the same genomic region can map to several sequences with near-equal scores. This reduces the standout score used for the mapping decision, causing some on-target reads to be discarded. Sigmoni’s *F*_1_ like-wise decreases as the reference collection grows for both the SARS-CoV-2 and *S. cerevisiae* classification tasks, with its *S. cerevisiae* recall slipping from 0.96 at *N* = 1 to 0.88 at *N* = 8. Its specificity stays near 1.0 because the off-target reference in its index allows it to filter out off-target reads.

**Fig. 2.**
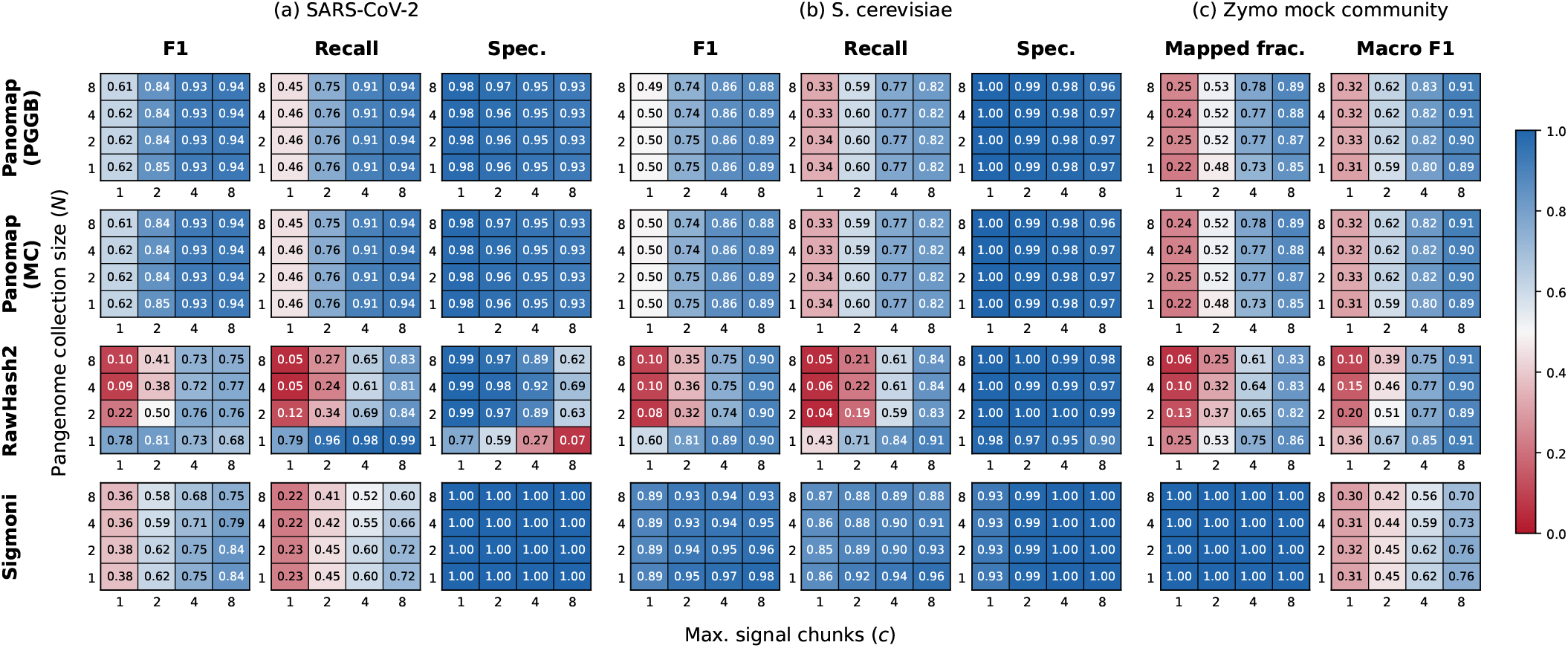
Cross-tool mapping accuracy across pangenome collection size *N* and maximum signal chunks *c* for (a) SARS-CoV-2, (b) *S. cerevisiae*, and (c) Zymo. Rows show tools and graph builders; columns show datasets and metrics.

Together, these results show that Panomap is robust to pangenome scaling when the reads are already well represented by a single reference.

### Improving accuracy through pangenome scaling when exact strains are absent

We next test whether adding related strains helps when the exact strain that produced the reads is absent from the reference collection. In this setting, the added strains diversify the reference collection and should help improve mapping.

We evaluate Panomap against RawHash2 and Sigmoni on a seven-way species classification task. We use the bacterial subset of the ZymoBIOMICS D6322 mock community reads curated by the Sigmoni authors, which includes 9,856 reads across seven bacterial species (yeast reads excluded) from [4]. Each tool assigns every read to one of the seven species. The reference includes all seven species, but represents each by related assemblies rather than the exact ZymoBIOMICS strain, mimicking a sample whose exact strains are unknown. For each species we scale the number of related assemblies from 1 to 8 (*N* = 1, 2, 4, 8). Ground-truth species labels come from basecalling each read and aligning it with minimap2 to per-species assemblies, following Sigmoni’s protocol. We report per-species *F*_1_ and summarize with macro *F*_1_, the mean across the seven species. Panomap maps to pggb and MC graphs, while RawHash2 and Sigmoni take the same assemblies as multi-FASTA reference collections.

From Figure 2c, Panomap’s accuracy improves as related assemblies are added: macro *F*_1_ rises from 0.89 at *N* = 1 to 0.91 at *N* = 8 at *c* = 8, with pggb and MC behaving the same. Conversely, RawHash2’s macro *F*_1_ at only one signal chunk drops from 0.36 at *N* = 1 to 0.10 at *N* = 8, though recovers to 0.91 at up to 8 signal chunks. Sigmoni does not gain from the added assemblies: its macro *F*_1_ stays below Panomap’s and slips from 0.76 to 0.70 as assemblies are added (at *c* = 8). Because Sigmoni always assigns a species label, it never leaves a read unmapped, whereas Panomap and RawHash2 can. Macro *F*_1_ reflects unmapped reads as misses, so it compares the three tools on equal footing despite their different mapped fractions.

Thus, when the exact strain is absent from the reference, Panomap benefits from the added diversity of related assemblies in the pangenome reference collection.

### Recovering reads from unrepresented divergent alleles at a polymorphic locus

Polymorphic loci can contain highly diverged alleles, making a single linear reference a poor representative of some sequenced samples. This is especially problematic when mapping decisions must be made from short signal prefixes. We test whether Panomap can recover reads from divergent alleles when the pangenome reference contains related allele groups but excludes the group that produced the reads.

We evaluate this setting at HLA-DRB1, a highly polymorphic locus. We use HLA-DRB1 alleles from the IPD-IMGT/HLA database [38], which are organized into 13 groups by sequence similarity. We use the GRCh38 HLA-DRB1 sequence, which is a group 15 allele [39], as the fixed single-reference baseline. We then select three source allele groups for read simulation based on their average divergence from this baseline: group 15 (0.08%), group 12 (27.29%), and group 7 (41.41%). These groups test cases where the simulated reads are close to, moderately diverged from, or far from the single-reference baseline. For each source allele group, we simulate 4,000 reads of ~8 kb mean length from alleles in that group using Squigulator with its R9.4.1 profile.

For each source allele group, we map the simulated reads against three references using Panomap. The first is the fixed GRCh38 HLA-DRB1 sequence (*N* = 1), which is the same single-reference baseline for all three source groups. The second is a small pangenome graph that excludes the source group and contains four alleles from each of the remaining 12 HLA-DRB1 groups (*N* = 48). The third is a larger pangenome graph that also excludes the source group and contains all alleles from the remaining 12 groups (*N* = max, 584–628 depending on the source group). Both pangenome references are built with pggb. Because neither pangenome includes the source group used to simulate the reads, any mapped reads must be recovered through related allele groups represented in the graph. We report recall, the fraction of simulated reads that Panomap maps, across *c* = 1, 2, 4, 8 maximum signal chunks.

As shown in Table 1, the benefit of the pangenome references depends on how far the read source group is from the GRCh38 baseline. For group 15 reads, the GRCh38 HLA-DRB1 sequence is itself a group 15 allele, so the fixed single-reference baseline already maps these reads well, with recall 0.936 at *c* = 1 and near 1.0 at *c* = 8. The two pangenome references exclude group 15, so they can only map these reads through related allele groups represented in the graph; their recall is slightly lower at *c* = 1 (0.908–0.911) but remains close to the baseline.

**Table 1.** HLA-DRB1 recall for reads from selected source allele groups. Reads are simulated from each selected group and mapped against the fixed GRCh38 HLA-DRB1 reference (*N* = 1) or pangenome references that exclude the source group (*N* = 48 or *N* = max).

| Read source group<br>(divergence %) | $N$ | Recall at maximum signal chunks $c$ | | | |
| --- | --- | --- | --- | --- | --- |
|  |  | 1 | 2 | 4 | 8 |
| Group 15<br>(0.08%) | 1 | 0.936 | 0.986 | 0.996 | 0.996 |
|  | 48 | 0.911 | 0.966 | 0.990 | 0.990 |
|  | 584 | 0.908 | 0.963 | 0.988 | 0.989 |
| Group 12<br>(27.29%) | 1 | 0.749 | 0.879 | 0.959 | 0.981 |
|  | 48 | 0.902 | 0.962 | 0.992 | 0.993 |
|  | 628 | 0.899 | 0.956 | 0.990 | 0.991 |
| Group 07<br>(41.41%) | 1 | 0.610 | 0.769 | 0.910 | 0.961 |
|  | 48 | 0.891 | 0.964 | 0.988 | 0.989 |
|  | 623 | 0.889 | 0.965 | 0.987 | 0.987 |

For group 12 reads, which are moderately diverged from GRCh38, the pangenome references improve recall most when the decision is made from few signal chunks. At *c* = 1, recall rises from 0.749 with GRCh38 to 0.902 with the *N* = 48 pangenome and 0.899 with the *N* = max pangenome. As *c* increases, the GRCh38 baseline catches up, reaching 0.981 at *c* = 8 compared with 0.993 and 0.991 for the two pangenome references.

For group 7 reads, which are farthest from GRCh38, the benefit is larger. At *c* = 1, recall rises from 0.610 with GRCh38 to 0.891 with the *N* = 48 pangenome and 0.889 with the *N* = max pangenome. The two pangenome references perform similarly, showing that the smaller 48-allele collection already captures most of the benefit. As *c* increases, GRCh38 again catches up, but still trails the pangenome references by a few percent at *c* = 8.

These results show that pangenome diversity improves recall when the fixed single-reference baseline is weakest: for source groups that are more diverged from GRCh38 and for decisions made from short signal prefixes. Panomap can therefore recover reads from polymorphic loci even when the allele group that produced the reads is not included in the pangenome reference.

### Index size and per-read processing time

We next examine the cost of using pangenome references. We measure index size and per-read processing time for Panomap, RawHash2, and Sigmoni on the three real-read benchmarks from the first two experiments: SARS-CoV-2, *S. cerevisiae*, and Zymo.

Figure 3 shows how each tool’s index size grows with the pangenome collection. Index size is measured as the on-disk byte size of the files each tool loads during mapping or classification (measurement commands and the files required by each tool are provided in the supplementary material). Panomap’s index scales sublinearly on all three datasets: as the collection grows eightfold from *N* = 1 to *N* = 8, the index stays near 1 MB on SARS-CoV-2 and grows only 2.2× on *S. cerevisiae* and 3.4× on Zymo. In contrast, RawHash2’s index grows near-linearly on all three datasets, reaching 3.17GB on Zymo, a 7.3× increase from *N* = 1 to *N* = 8. This is because RawHash2 treats each reference sequence as a separate target and stores sequences shared across reference sequences redundantly. Sigmoni’s index also scales sublinearly and is the smallest of the three on *S. cerevisiae* and Zymo, growing 1.65× and 2.53× from *N* = 1 to *N* = 8 (to 111 MB and 329 MB). On SARS-CoV-2, however, its index is the largest of the three (17.4 MB), because Sigmoni’s binary classification mode also requires an off-target reference, and the *E. coli* null dominates the small SARS-CoV-2 target collection.

**Fig. 3.**
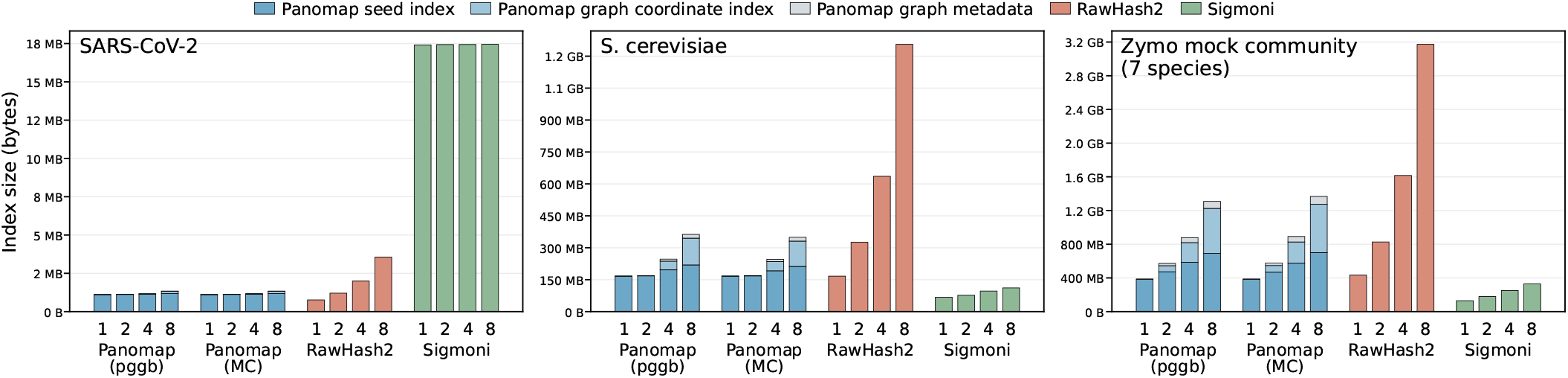
Index size scaling across pangenome collection size *N* for Panomap, RawHash2, and Sigmoni. The Panomap breakdown separates the seed index, graph coordinate index, and other graph metadata.

Panomap’s sublinear index size scaling comes from storing seeds for shared graph nodes once rather than redundantly per path. Its other components, the graph coordinate index and graph metadata, grow with the graph itself, so the total index size follows the graph’s node count. This differs sharply across datasets: SARS-CoV-2’s graph stays under 2,000 nodes even at *N* = 8, whereas Zymo’s reaches 5.7– 6.3 M nodes (Table S5). Sigmoni’s compact index comes with a separate cost: because its binary classification mode requires an off-target reference, the total index size also depends on that reference. In our SARS-CoV-2 experiment, the *E. coli* off-target reference already dominates the index. If the off-target reads were human and hg38 were used as the offtarget reference, Sigmoni’s index would expand to 14.5 GB.

We measure per-read processing time as total wall-clock time divided by the number of reads, using 64 threads (Table S8). Panomap is competitive overall. It is fastest on SARS-CoV-2, staying below 0.25 ms per read for every tested combination of *N* and *c*, and on *S. cerevisiae* all three tools converge to 2–3 ms per read at *N* = 8 and *c* = 8. On Zymo, however, Panomap is slowest when *N* = 8 and *c* = 8, taking 4.7 ms per read compared with 2.5 ms for RawHash2 and 4.4 ms for Sigmoni. This slowdown reflects the cost of Panomap’s current mapping approach as the pangenome grows. In the current design, graph anchors are expanded onto paths and chained independently within each path, so chaining work grows with the number of paths represented in the graph. On *S. cerevisiae* reads at *c* = 8, Panomap’s per-read time grows from 0.83 ms at *N* = 1 to 3.0 ms at *N* = 8. Chaining directly on the graph could reduce this dependence on the number of paths, but would require robust graph-distance methods.

Overall, Panomap keeps index growth sublinear by sharing seeds across graph nodes, while retaining competitive per-read time across the real-read benchmarks. Its main cost is increased read mapping time from per-path chaining, making chaining directly on graph anchors the clearest target for future optimization.

## Discussion

### Chaining on Pangenome Graphs in Signal Space

Colinear chaining relies on evaluating distance between anchors on both the read and the reference. While this is straightforward on a linear reference, graph references introduce ambiguity because multiple paths and cycles may connect the same pair of anchors. Base-space sequence-to-graph mappers address this challenge using graph-level distance metrics, including shortest-path (minigraph [23]), snarl-based (vg giraffe [21]), and topological (GraphAligner [22]) formulations. In practice, these metrics are often sufficient because chaining primarily identifies candidate regions, while a downstream alignment stage, such as partial-order alignment used in vg map [19, 40] or bit-parallel sequence-to-graph alignment [41] used in GraphAligner, recovers the exact read-to-graph correspondence.

Signal-space mapping currently lacks an analogous correction stage. Although sequence-to-graph alignment has matured substantially, signal-to-graph alignment remains computationally challenging. Dynamic programming formulations for signal-to-graph alignment [42] can be derived by extending sequence-to-graph alignment algorithms to dynamic time warping, but the accelerations that make practical sequence-to-graph aligners fast, such as bit-parallel scoring over binary match/mismatch, do not transfer easily to the dynamic-time-warping setting where signal distances are continuous. Consequently, errors introduced during chaining are more difficult to recover downstream, placing greater importance on accurate graph distance estimates that reflect nucleotide distance along paths.

Panomap addresses this challenge by chaining anchors within each path. Rather than approximating distances directly on the graph, Panomap projects graph anchors onto the paths that contain them and chains the projected anchors within each path’s coordinate system. This preserves path-specific nucleotide distances and also makes repeated traversals through cycles explicit. Across our evaluations, this strategy preserved accuracy as pangenome collections scaled when the single-reference baseline maps well and improved recall on absent target alleles in a polymorphic region. These results suggest that chaining with exact distances along paths is sufficient to extend pangenome graph mapping into signal space while retaining the benefits of pangenome representation with variation graphs.

Future work may move beyond per-path chaining through more accurate graph distance metrics, graph-aware chaining formulations, or practical signal-to-graph alignment algorithms. Progress in any of these directions could improve scalability and facilitate signal-space mapping on increasingly large pangenomes, including human-scale variation graphs [43].

### Signal Representations and Early Decision Making

Beyond graph-specific challenges, our experience developing Panomap suggests that signal representation quality remains a dominant challenge across signal-space methods. Event segmentation, tokenization, and seed generation each introduce opportunities for divergence between reference-derived and read-derived representations, ultimately affecting the mapper’s ability to extract reliable matches from short signal prefixes.

Event segmentation and tokenization are two major sources of this representation error. Many signal-space tools rely on the t-statistic detector introduced by Scrappie [35] to partition the continuous nanopore signals into discrete events. However, segmentation quality can vary with detector parameters, sequencing chemistry, and signal characteristics. Over-segmentation may split one expected event into multiple detected events, while under-segmentation may merge distinct events together. Tokenization introduces an additional source of error, where similar events may be assigned different tokens when they fall near quantization boundaries, or distinct events may collapse into the same token when the alphabet is too coarse. Both cases reduce the reliability of subsequent exact seed matching. As a result, signal-space methods often require denser seeding than their base-space counterparts, using shorter seeds or less aggressive seed subsampling. This increases seed frequency, index size, candidate anchor counts, and chaining cost.

These representation errors affect both final mapping accuracy and early decision making. Fewer reliable seeds produce weaker chains and more ambiguous candidate mappings. This is particularly important in adaptive sampling, where decisions must be made from short signal prefixes. Reads that cannot be confidently classified from the current signal chunks must continue sequencing so that Panomap can collect more signal and attempt another decision. Thus, although signal-space methods avoid the computational cost of basecalling, representation errors can leave their recall on the earliest chunks below that of base-space adaptive sampling such as [5], which, when a GPU basecaller is available, can already make reliable decisions from a short signal prefix.

Signal-space methods remain useful as adaptive samplers in resource-constrained environments, and as pre-basecall filters running alongside sequencing to identify confident cases early and reduce downstream basecalling load [25, 44].

Future improvements in event detection, tokenization, seed design, or learned signal representations could improve mapping quality, reduce index and candidate-set sizes. By making decisions more reliable from shorter read prefixes, improved representations would make signal-space adaptive sampling more competitive with base-space approaches while preserving its key advantage of avoiding basecalling for early decisions.

## Conclusion

Panomap is the first signal-space mapper that operates on pangenome variation graphs. Panomap preserves mapping accuracy when a single reference already represents the sample well, improves mapping when related pangenome diversity better represents a sample whose exact strain is absent, and improves recall at a polymorphic locus when the sequenced allele is missing from the reference. By representing shared sequences once while preserving the paths in the graph, Panomap brings population-aware reference representation into signal-space mapping. More broadly, our results identify graph-aware chaining and robust signal representation as key directions for making signal-space mapping more accurate, scalable, and robust to genetic diversity.

## Data Availability

Panomap source code is available at https://github.com/cornell-brg/panomap. The datasets and references used in this study were curated from the accessions and sources cited in the Methods and Supplementary Material and are available directly on Zenodo at doi:10.5281/zenodo.21420010 [45].

## AI Use Acknowledgement

The authors used Claude Code and ChatGPT as coding and editorial aids under author supervision. Full details are provided in the Supplementary Information.

## Supplementary Material

### S1. Read datasets

The three real-read datasets used in the evaluation use R9.4.1 chemistry at 4 kHz. For the SARS-CoV-2 and *S. cerevisiae* datasets we take the first 40,000 reads from each source, converted from FAST5 to BLOW5 with `slow5tools f2s`. The Zymo reads dataset is the bacterial portion of the set curated by the authors of Sigmoni with yeast reads excluded.

**Table 1.** Datasets used in cross-tool evaluations.

| Dataset | Chemistry | Reads used | Data source |
| --- | --- | --- | --- |
| SARS-CoV-2 | R9.4.1 | 40,000 | InterARTIC SARS-CoV-2 (seq.bioinf.science/interartic-corona) |
| *S. cerevisiae* | R9.4.1 | 40,000 | SRA SRR8648503 (BioProject PRJNA510813) |
| Zymo D6322 | R9.4.1 | 9,856 | subset of SRA SRX7711546 curated by the Sigmoni authors; yeast reads excluded |

The binary classification task in the evaluation uses *E. coli* reads simulated using Squigulator (v0.5.0) with K-12 MG1655 (NCBI `NC_000913.3`). Command used to simulate reads is:

~~~
squigulator -x dna-r9-min --seed 42 \
-r 8000 -n 40000 --sample-rate 4000 \
-o ecoli-sim-r9-4khz-40k-8kb.blow5 \
-q ecoli-sim-r9-4khz-40k-8kb.fasta \
-a ecoli-sim-r9-4khz-40k-8kb.paf \
ecoli_MG1655.fa
~~~

For the HLA-DRB1 experiment, we simulate 4,000 reads from each of groups 15, 12, and 7 using:

~~~
squigulator -x dna-r9-min --seed 42 \
-r 8000 -n 4000 --sample-rate 4000 \
-o <drb1-group>.blow5 \
-q <drb1-group>.fa \
-a <drb1-group>.paf \
<drb1-group-alleles>.fa
~~~

The reported divergence for group 15 (0.08%), group 12 (27.29%), and group 7 (41.41%) is the average divergence of each group’s alleles from the GRCh38 DRB1 allele, *DRB1*15:01:01:01*. We compute this by aligning each allele to the GRCh38 allele with `minimap2 -x asm20 -c -secondary=no` and averaging `NM` divided by aligned block length over the best primary alignments.

### S2. Reference collections

For the binary classification (SARS-CoV-2 and *S. cerevisiae* and multi-class classification (Zymo) experiments that vary the reference collection size, we build each collection by taking the first *N* references listed for that organism in Tab. S2. Thus, *N* = 1 uses Ref 1, *N* = 2 uses Ref 1–Ref 2, *N* = 4 uses Ref 1–Ref 4, and *N* = 8 uses Ref 1–Ref 8. We use this construction for SARS-CoV-2, *S. cerevisiae*, and the Zymo species. The *S. cerevisiae* references come from the pangenome panel available under ENA BioProject `PRJEB7245`. The other references are listed by accession in the table.

**Table 2.** Per-organism reference pool. Each cell shows the strain or lineage name on the top line and the corresponding accession on the bottom line. Reference collections of size *N* use the first *N* entries in each row.

| Organism | Ref 1 | Ref 2 | Ref 3 | Ref 4 | Ref 5 | Ref 6 | Ref 7 | Ref 8 |
| --- | --- | --- | --- | --- | --- | --- | --- | --- |
| SARS-CoV-2 | Wuhan-Hu-1<br>NC_045512.2 | B.1.1.7<br>OZ303776.1 | B.1.351<br>MW598419.1 | P.1<br>MZ264787.1 | B.1.617.2<br>PX952986.1 | BA.1<br>PX503961.1 | BA.5<br>PX497191.1 | XBB.1.5<br>PQ017954.1 |
| *S. cerevisiae* | SGDref<br>-- | S288C<br>-- | DBVPG6044<br>-- | DBVPG6765<br>-- | SK1<br>-- | UWOPS034614<br>-- | Y12<br>-- | YPS128<br>-- |
| *E. coli* | MG1655<br>NC_000913.3 | Sakai<br>NC_002695.2 | ATCC 25922<br>CP009072.1 | CFT073<br>NC_004431.1 | SE11<br>NC_011415.1 | BL21(DE3)<br>CP053602.1 | W<br>CP002967.1 | ED1a<br>NC_011745.1 |
| *B. subtilis* | 168<br>NC_000964.3 | BSn5<br>CP010053.1 | NCIB3610<br>CP002468.1 | CF03<br>NZ_CP110105.1 | HC-9<br>NZ_CP149444.1 | LTH 7164<br>NZ_CP160208.1 | PY79<br>NZ_CP196824.1 | 6051-HGW<br>CP003329.1 |
| *E. faecalis* | 62<br>NC_017316.1 | DENG1<br>NC_018221.1 | RF_1_1_25_1<br>NZ_CP046247.1 | L13<br>NZ_CP071172.1 | W5<br>NZ_CP118755.1 | SVJ-EF01<br>NZ_CP121233.1 | OG1RF<br>CP002621.1 | V583<br>NC_004668.1 |
| *L. monocytogenes* | FSL R2-561<br>NC_017547.1 | F2365<br>NC_002973.6 | EGD-e<br>NC_003210.1 | HCC23<br>NC_011660.1 | J1-220<br>NC_018587.1 | J1816<br>NC_018584.1 | M7<br>NC_018589.1 | N1-011A<br>NC_021830.1 |
| *P. aeruginosa* | DSM 50071<br>CP004061.1 | LESB58<br>NC_011770.1 | M18<br>CP008857.1 | PAO1-UW<br>NZ_CP079712.1 | 2-216<br>NZ_CP158049.1 | PB2<br>NZ_CP172964.1 | M25569<br>NZ_CP197382.1 | PAO1<br>NC_002516.2 |
| *S. enterica* | Choleraesuis<br>NC_006905.1 | Enteritidis<br>NC_011294.1 | Heidelberg<br>NC_011083.1 | Corvallis<br>NZ_CP175687.1 | Derby<br>NZ_CP175995.1 | Paratyphi A<br>NC_006511.1 | Newport<br>NC_011080.1 | Typhimurium<br>NC_003197.2 |
| *S. aureus* | COL<br>NC_002951.2 | JH9<br>NC_009487.1 | MRSA252<br>NC_002952.2 | 11819-97<br>NC_017351.1 | N315<br>NC_002758.2 | Mu50<br>NC_002745.2 | NCTC 8325<br>NC_007795.1 | USA300<br>NC_010079.1 |

### S3. Graph construction

Pangenome graphs are built with pggb (v0.7.2) and Minigraph-Cactus (v3.1.4).

pggb command:

~~~
pggb -i <input>.fa.gz -n <N> -o <out-dir> -t 16 -p <P> -s <S>
~~~

Per-dataset settings:

**Table 3.** pggb parameters used per dataset.

| Dataset | $N$ | $P$ (%) | $S$ (bp) |
| --- | --- | --- | --- |
| SARS-CoV-2 | 2, 4, 8 | 90 | 1000 |
| *S. cerevisiae* | 2, 4, 8 | 90 | 1000 |
| Zymo species* | 2, 4, 8 | 90 | 5000 |
| HLA-DRB1 | 48, max | 90 | 1000 |

\* Per-species GFAs are merged into one multi-component GFA with `concat_gfa_components.py`, which shifts each species' segment IDs by a per-species offset and concatenates the S, L, and P lines.

Minigraph-Cactus command:

~~~
cactus-pangenome jobstore-<N> <input>.seqfile \
--outDir mc-build-<N> --outName <name>-mc-<N> \
--reference <ref> --gfa clip \
--mgCores 16 --maxCores 16 \
--workDir work-<N> --clean always \
--batchSystem single_machine
~~~

Per-dataset settings:

**Table 4.** Minigraph-Cactus parameters used per dataset.

| Dataset | $N$ | Reference |
| --- | --- | --- |
| SARS-CoV-2 | 2, 4, 8 | wuhan |
| *S. cerevisiae* | 2, 4, 8 | SGDref |
| Zymo species* | 2, 4, 8 | first alphabetical strain per species |

\* Per-species GFAs are merged into one multi-component GFA with `concat_gfa_components.py` (same as for `pggb`).

For single haplotype references (*N* = 1), no graph builder is run: the single reference FASTA is converted to GFA format with one segment (*S*) line and one path (*P*) line per contig. This is done with `fa2gfa.py`.

The resulting graph statistics (node, edge, and path counts as stored in the Panomap index) are listed in Tab. S5.

**Table 5.** Pangenome graph statistics per (dataset, builder, *N*). Node, edge, and path counts are reported as stored in the Panomap index (equal to 2× the GFA segment / link / path count due to the canonical forward / reverse-complement node-pair convention).

| Dataset | Builder | $N$ | Nodes | Edges | Paths |
| --- | --- | --- | --- | --- | --- |
| SARS-CoV-2 | pggb | 1 | 2 | 0 | 2 |
| SARS-CoV-2 | pggb | 2 | 154 | 202 | 4 |
| SARS-CoV-2 | pggb | 4 | 454 | 604 | 8 |
| SARS-CoV-2 | pggb | 8 | 1,420 | 1,912 | 16 |
| SARS-CoV-2 | MC | 1 | 2 | 0 | 2 |
| SARS-CoV-2 | MC | 2 | 194 | 242 | 4 |
| SARS-CoV-2 | MC | 4 | 488 | 642 | 8 |
| SARS-CoV-2 | MC | 8 | 1,420 | 1,912 | 16 |
| *S. cerevisiae* | pggb | 1 | 34 | 0 | 34 |
| *S. cerevisiae* | pggb | 2 | 4,372 | 6,004 | 68 |
| *S. cerevisiae* | pggb | 4 | 699,852 | 942,296 | 136 |
| *S. cerevisiae* | pggb | 8 | 1,279,144 | 1,732,748 | 272 |
| *S. cerevisiae* | MC | 1 | 34 | 0 | 34 |
| *S. cerevisiae* | MC | 2 | 27,934 | 29,568 | 72 |
| *S. cerevisiae* | MC | 4 | 752,488 | 1,011,798 | 148 |
| *S. cerevisiae* | MC | 8 | 1,288,810 | 1,740,706 | 296 |
| Zymo | pggb | 1 | 14 | 0 | 14 |
| Zymo | pggb | 2 | 1,949,234 | 2,603,208 | 28 |
| Zymo | pggb | 4 | 4,099,566 | 5,500,282 | 56 |
| Zymo | pggb | 8 | 5,679,886 | 7,635,388 | 112 |
| Zymo | MC | 1 | 14 | 0 | 14 |
| Zymo | MC | 2 | 2,128,924 | 2,835,456 | 34 |
| Zymo | MC | 4 | 4,485,834 | 6,015,228 | 110 |
| Zymo | MC | 8 | 6,279,702 | 8,440,784 | 180 |

### S4. Running the tools

All tools use 16 threads for indexing and 64 threads for mapping. Versions: panomap (commit `af79d3`), RawHash2 v2.2, Sigmoni (commit `c4f220b`) backed by SPUMONI v2.0.7.

panomap index command:

~~~
panomap index <gfa> -m r9.4 -t 16 -o <out>.pirx
~~~

panomap map command:

~~~
panomap map --index <idx> --chainer path-chain <map-args> \
--max-chunks <c> -t 64 <reads>.blow5 -o <out>.gaf
~~~

Per-dataset `<map-args>`:

**Table 6.**
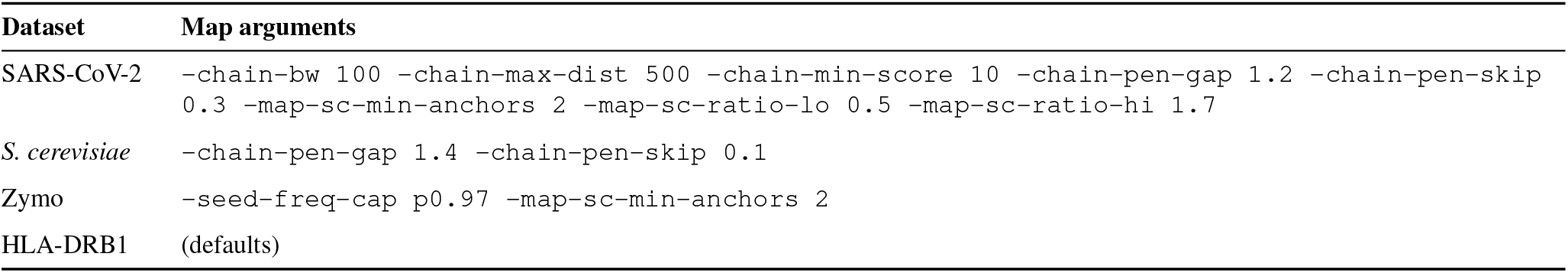
panomap per-dataset map arguments.

| Dataset | Map arguments |
| --- | --- |
| SARS-CoV-2 | -chain-bw 100 -chain-max-dist 500 -chain-min-score 10 -chain-pen-gap 1.2 -chain-pen-skip 0.3 -map-sc-min-anchors 2 -map-sc-ratio-lo 0.5 -map-sc-ratio-hi 1.7 |
| *S. cerevisiae* | -chain-pen-gap 1.4 -chain-pen-skip 0.1 |
| Zymo | -seed-freq-cap p0.97 -map-sc-min-anchors 2 |
| HLA-DRB1 | (defaults) |

RawHash2 index command:

~~~
rawhash2 -x <preset> -t 16 -p r9.4.model -d <out>.ind <ref>.fa
~~~

RawHash2 map command:

~~~
rawhash2 -x <preset> --max-chunks <c> -t 64 \
-o <out>.paf <idx> <reads>.blow5
~~~

Per-dataset `<preset>`:

**Table 7.** RawHash2 per-dataset preset.

| Dataset | Preset |
| --- | --- |
| SARS-CoV-2 | viral |
| *S. cerevisiae* | sensitive |
| Zymo | fast |

Sigmoni index command:

~~~
sigmoni-index --shred 100000 -p <pos.fa> -n <null.fa> \
--spumoni-path <spumoni> -o <out-dir> --ref-prefix <name>
~~~

Sigmoni map command:

~~~
sigmoni --thresh 1.0 --sp --complexity \
-i <reads>.blow5 -r <idx-prefix> -o <out-dir> \
--max-chunks <c> -t 64
~~~

All Sigmoni runs use `-thresh 1.0` and `-shred 100000`.

### S5. Index size measurement

Index size was measured as the on-disk byte size of the files each tool loads during mapping or classification, using `du -sb`. For Panomap and RawHash2, this is the single serialized index file loaded at mapping time: `.pirx` for Panomap and `.ind` for RawHash2. For Sigmoni, we counted only the files loaded at classification time: the r-index (`*.thrbv.spumoni`), the reference document array (`*.doc`), the empirical null database (`*.pmlnulldb`, present in binary mode only), and the document-to-class file lists (`filelist.txt` for multi-class classification, or `pos_filelist.txt` and `null_filelist.txt` for binary classification). We excluded build-time artifacts that are not read during classification, including the binned reference and per-shred FASTA files, `spumoni_null_reads.fa`, `poremodel.bins`, `*.fdi`, and the statistics log.

### S6. Per-read processing time

Per-read processing time was computed as elapsed wall-clock time divided by the number of reads and converted to milliseconds. Elapsed wall-clock time was measured with GNU `/usr/bin/time` using `%e`. All tools were run with 64 threads on the same host used for the main experiments. Results are reported across pangenome collection size *N* and maximum signal chunks *c*.

**Table 8.** Per-read processing time (ms) across datasets, tools, *N*, and *c*.

| Dataset | Tool | $N$ | $c = 1$ | $c = 2$ | $c = 4$ | $c = 8$ |
| --- | --- | --- | --- | --- | --- | --- |
| SARS-CoV-2 | Panomap (pggb) | 1 | 0.23 | 0.23 | 0.23 | 0.23 |
|  |  | 2 | 0.23 | 0.23 | 0.23 | 0.23 |
|  |  | 4 | 0.23 | 0.23 | 0.24 | 0.24 |
|  |  | 8 | 0.23 | 0.23 | 0.24 | 0.24 |
|  | Panomap (MC) | 1 | 0.23 | 0.24 | 0.24 | 0.24 |
|  |  | 2 | 0.23 | 0.24 | 0.24 | 0.24 |
|  |  | 4 | 0.23 | 0.24 | 0.24 | 0.24 |
|  |  | 8 | 0.23 | 0.24 | 0.24 | 0.24 |
|  | RawHash2 | 1 | 0.30 | 0.30 | 0.30 | 0.30 |
|  |  | 2 | 0.30 | 0.30 | 0.32 | 0.34 |
|  |  | 4 | 0.30 | 0.31 | 0.36 | 0.40 |
|  |  | 8 | 0.30 | 0.33 | 0.43 | 0.50 |
|  | Sigmoni | 1 | 0.59 | 0.76 | 1.06 | 1.55 |
|  |  | 2 | 0.60 | 0.75 | 1.03 | 1.53 |
|  |  | 4 | 0.60 | 0.75 | 1.06 | 1.52 |
|  |  | 8 | 0.61 | 0.75 | 1.05 | 1.49 |
| *S. cerevisiae* | Panomap (pggb) | 1 | 0.65 | 0.69 | 0.75 | 0.83 |
|  |  | 2 | 0.69 | 0.78 | 0.85 | 1.02 |
|  |  | 4 | 0.83 | 1.01 | 1.28 | 1.72 |
|  |  | 8 | 1.08 | 1.48 | 2.05 | 3.04 |
|  | Panomap (MC) | 1 | 0.65 | 0.69 | 0.75 | 0.84 |
|  |  | 2 | 0.70 | 0.78 | 0.86 | 1.04 |
|  |  | 4 | 0.84 | 1.02 | 1.27 | 1.71 |
|  |  | 8 | 1.04 | 1.42 | 1.94 | 2.85 |
|  | RawHash2 | 1 | 0.71 | 0.72 | 0.77 | 0.79 |
|  |  | 2 | 0.73 | 0.76 | 0.83 | 0.90 |
|  |  | 4 | 0.80 | 0.84 | 1.08 | 1.29 |
|  |  | 8 | 0.92 | 1.06 | 1.71 | 2.51 |
|  | Sigmoni | 1 | 1.13 | 1.34 | 1.75 | 2.43 |
|  |  | 2 | 1.18 | 1.38 | 1.75 | 2.44 |
|  |  | 4 | 1.21 | 1.44 | 1.84 | 2.52 |
|  |  | 8 | 1.35 | 1.56 | 1.92 | 2.66 |
| Zymo | Panomap (pggb) | 1 | 1.53 | 1.55 | 1.60 | 1.70 |
|  |  | 2 | 1.68 | 1.80 | 1.99 | 2.24 |
|  |  | 4 | 1.97 | 2.27 | 2.68 | 3.29 |
|  |  | 8 | 2.38 | 2.86 | 3.67 | 4.70 |
|  | Panomap (MC) | 1 | 1.51 | 1.55 | 1.62 | 1.71 |
|  |  | 2 | 1.65 | 1.80 | 2.02 | 2.27 |
|  |  | 4 | 1.88 | 2.29 | 2.64 | 3.29 |
|  |  | 8 | 2.50 | 2.93 | 3.72 | 4.85 |
|  | RawHash2 | 1 | 1.80 | 1.78 | 1.78 | 1.79 |
|  |  | 2 | 1.99 | 1.83 | 1.85 | 1.87 |
|  |  | 4 | 1.97 | 1.88 | 1.90 | 1.94 |
|  |  | 8 | 2.34 | 2.00 | 2.34 | 2.46 |
|  | Sigmoni | 1 | 2.67 | 2.82 | 3.13 | 4.05 |
|  |  | 2 | 2.62 | 2.94 | 3.27 | 4.12 |
|  |  | 4 | 2.78 | 3.05 | 3.40 | 4.11 |
|  |  | 8 | 2.96 | 3.24 | 3.55 | 4.40 |

### S7. AI Use Disclosure

The authors used Claude Code and ChatGPT in the following categories, under author guidance:

- *Text refinement*. Prose drafts written by the human authors were edited and polished for clarity with AI suggestions. Suggested rewrites were reviewed, accepted, modified, or rejected by the authors.
- *Code assistance, Panomap software development*. Claude Code was used heavily during the implementation of the Panomap mapper as a coding aid under author guidance. All algorithmic design, mapping semantics, data structures, and final correctness of the implementation are the responsibility of the authors.
- *Code assistance, figures*. Figure-generation Python scripts were drafted and refined for layout, font sizing, color palette, and similar visual elements with AI assistance. All data, numerical results, and figure values were produced by the authors’ analysis pipelines.
- *Code assistance, analysis and experiments*. Experiment runner scripts, data-parsing scripts, and TSV format conversions were drafted with AI assistance. All scripts were reviewed and verified by the authors before use; all reported numerical results were produced by these reviewed scripts.
- *Table formatting*. L^A^T_E_X table structure and layout (column alignment, multirow grouping, spacing) was refined with AI assistance.

